# Extensive tonotopic mapping across auditory cortex is recapitulated by spectrally-directed attention, and systematically related to cortical myeloarchitecture

**DOI:** 10.1101/142513

**Authors:** Frederic K. Dick, Matt I. Lehet, Martina F. Callaghan, Tim A. Keller, Martin I. Sereno, Lori L. Holt

## Abstract

Auditory selective attention is vital in natural soundscapes. But, it is unclear how attentional focus on the primary dimension of auditory representation - acoustic frequency - might modulate basic auditory functional topography during active listening. In contrast to visual selective attention, which is supported by motor-mediated optimization of input across saccades and pupil dilation, the primate auditory system has fewer means of differentially sampling the world. This makes spectrally-directed endogenous attention a particularly crucial aspect of auditory attention. Using a novel functional paradigm combined with quantitative MRI, we establish that human frequency-band-selective attention drives activation in both myeloarchitectonically-estimated auditory core, and across the majority of tonotopically-mapped non-primary auditory cortex. The attentionally-driven best-frequency maps show strong concordance with sensory-driven maps in the same subjects across much of the temporal plane, with poor concordance in non-auditory areas. There is significantly greater activation across most of auditory cortex when best frequency is attended, versus ignored. Moreover, the single frequency bands that evoke the least activation and the frequency bands that elicit the least activation when attention is directed to them also correspond closely. Finally, the results demonstrate that there is spatial correspondence between the degree of myelination and the strength of the tonotopic signal across a number of regions in auditory cortex. Strong frequency preferences across tonotopically-mapped auditory cortex spatially correlate with R_1_-estimated myeloarchitecture, indicating shared functional and anatomical organization that may underlie intrinsic auditory regionalization.

**Significance:** Perception is an active process especially sensitive to attentional state. Listeners direct auditory attention to track a violin’s melody within an ensemble performance, or to follow a voice in a crowded cafe. Although diverse pathologies reduce quality of life by impacting such spectrally-directed auditory attention, its neurobiological bases are unclear. We demonstrate that human primary and non-primary auditory cortical activation is modulated by spectrally-directed attention in a manner that recapitulates its tonotopic sensory organization. Further, the graded activation profiles evoked by single frequency bands are correlated with attentionally-driven activation when these bands are presented in complex soundscapes. Finally, we observe a strong concordance in the degree of cortical myelination and the strength of tonotopic activation across several auditory cortical regions.

## Introduction

Listeners shift attention across multiple simultaneously-present acoustic dimensions to home in on those that are diagnostic in guiding behavior (Idemaru and Holt, 2011; Herrmann and Henry, 2013; Shamma and Fritz, 2014). In nonhuman animal studies task-based spectral attention adaptively modulates auditory neurons’ spectrotemporal response fields (Fritz et al., 2010). Human neuroimaging results reveal that attention to streams of high-versus low-frequency acoustic input can modulate activity in tonotopically-defined regions (Paltoglou et al., 2009), as can imagery that evokes higher versus lower frequencies (Oh et al., 2013). In and directly around Heschl’s gyrus, there are strong frequency-band-specific attentional effects to high and low pure-tone streams presented to opposite ears (Da Costa et al., 2013) and a shared topography of sensory and attentionally-driven responses (Riecke et al., 2016). These results establish that endogenous attention directed across acoustic frequency, the primary axis of auditory representation, can modulate human cortical activity in a tonotopic manner around Heschl’s gyrus. But, there remain important unanswered questions about the neurobiological basis of human spectrally-directed attention.

First, does human primary auditory cortex exhibit attentionally-driven tonotopic organization? Non-human animal physiology establishes spectrally-directed attention in myelo-and cyto-architectonically-defined primary areas in ‘auditory core’ (Fritz et al., 2007b; Shamma and Fritz, 2014). However, although two recent neuroimaging studies have shown strong similarities between stimulus- and attentionally-driven tonotopic organization in and directly around Heschl’s gyrus (Da Costa et al., 2013; Riecke et al., 2016), it has not yet been possible to unambiguously localize this effect to human auditory core. Here, we use high-resolution quantitative MRI to estimate myelo-architectonically-defined auditory core, and demonstrate that spectrally-directed attention modulates its activation in a tonotopically-organized manner.

Second, is attentionally-driven tonotopic organization present outside of auditory core? In humans, (Riecke et al., 2016) found no significant evidence for tonotopically organized effects of spectral attention outside of early auditory areas, but did show that the information content of non-primary cortical frequency representations was sufficient for above-chance decoding of listeners’ frequency-selective attentional focus. The lack of attentionally-driven tonotopic mapping contrasts with the finding that most non-primary cortical visual areas exhibit strong retinotopically-specific attentional effects (Saygin and Sereno, 2008)). Using intensive data collection (>7000 functional volumes per subject) we present evidence for widespread, tonotopically-organized modulation by spectral attention across much of auditory cortex, with individual differences in individual participants’ tonotopic maps reproduced in attentionally-driven maps.

Third, what is the effect of when frequency-selective attention is directed to a voxel’s nonpreferred frequency band? Detailed fMRI studies of stimulus-driven frequency response functions (Schönwiesner and Zatorre, 2009; Moerel et al., 2013) have shown graded and multipeaked frequency responses across human auditory cortex. However, it is unclear whether these more complex patterns are recapitulated by attention to a given frequency band. In the context of three distinct frequency bands, (Riecke et al., 2016) found that attentional filters appeared to be bandpass in and around Heschl’s gyrus. Here, using a five-frequency-band paradigm, we establish that graded response profiles evoked by single frequency bands are strongly associated with attentionally-driven response profiles to those frequencies across much of auditory cortex. We also show that a systematic topography of ‘dis-preferred’ frequency can be driven by attention, and establish the regionalization of spectral attentional effects relative to prior studies of crossmodal auditory attention (Petkov et al., 2004).

Finally, is there spatial correspondence between auditory cortical anatomy, as measured by the local change in R_1_-estimated myelination, and the strength of the fMRI-assessed strength of relative frequency selectivity? Post-mortem Gallyas staining to establish human myeloarchitecture reveals considerable variability in auditory cortical myelination that is associated with MRI signal change in the same brain (Wallace et al., 2016). Likewise, variation in cortical myelination estimated using T1/T2 ratio approaches also appears to correspond spatially with some functional variation in the superior temporal lobe (Glasser et al., 2016). Here, we demonstrate that there is spatial concordance between the degree of myelination and the amplitude of the frequency-selective tonotopic signal across several regions in auditory cortex.

## Methods

### Experiment Overview

We used a novel paradigm in which listeners direct attention to a series of four-tone ‘mini-sequences’ that fall within one of five possible spectral bands, without any spatial cues. The task is to monitor for temporally-adjacent mini-sequence repeats within the attended band. Inasmuch as this places a very high demand on encoding and integrating spectral sequences within a delimited frequency range, we expect it to be especially effective in evoking strong responses in non-primary auditory cortical areas. The goal is to address where, specifically, in the auditory system spectral gain from attention is evident, and akin to longstanding work in vision (Kastner and Ungerleider, 2000), to delineate the topographic maps across which attentional modulation is apparent.

The target mini-sequences were embedded in either an informationally-sparse or informationally-dense acoustic scene (Figure (Fig.) 1). Streams of four-tone mini-sequences were presented in either a single band [‘tonotopy’, Fig. 1a], or accompanied by mini-sequences in a ‘distractor’ frequency band, the center frequency of which varied in the frequency distance from the attended band across blocks [‘attention-tonotopy’, abbreviated ‘attn-tono’, Fig. 1b]. A verbal cue directed listeners’ attention to a specific frequency band, within which listeners monitored four-tone mini-sequences for repeats; the ‘distractor’ band in attn-tono blocks also contained repeats. Using a discretized version of a phase-encoded fMRI design (Sereno et al., 1995; Schwarzkopf et al., 2011; Rao:2005fn; Herdener et al., 2013; Langers et al., 2014), the cued frequency band stepped up or down in orderly steps across the acoustic spectrum across a 64-sec cycle [Fig. 1c]. Phase-encoded tonotopic designs benefit from the power and robustness of the ‘travelling wave’ method for topographic cortical mapping (Engel, 2012); the discretized (blocked) version we use here has the advantage of being able to be analyzed using both Fourier and regression approaches. This allowed us to include an additional, ‘randomized’ attn-tono condition that contributed both as a control condition in Fourier analyses, and also as an additional attn-tono run in regression analyses (see Fig.1d). The tone stimuli from this condition were *identical* to the ‘stepped’ attn-tono condition, but the order of the verbal cues directing listeners’ attention to a specific frequency band was scrambled in their assignment to blocks. This preserved the acoustics across conditions, but eliminated the consistent ‘stepping’ of attention through the frequency spectrum in the randomized condition thereby destroying the consistent phase-lag associated with specific frequency bands that support Fourier analyses [schematized in Fig. 1e, and see below]. Thus, to the extent that there are attentionally-driven frequency-selective maps in auditory cortex we expect tonotopically-organized attentional maps to be apparent in the ‘stepped,’ but not the ‘randomized’ attention-o-tonotopy conditions under Fourier analyses. In contrast, regression analyses include a model of attention, allowing ‘stepped’ and ‘randomized’ attn-tono conditions to be pooled to investigate the impact of attention on cortical activation. Across both Fourier and regression analyses, the ‘stepped’ attn-tono conditions were collapsed across runs for which the cued frequency band stepped up in frequency and those that stepped down; inclusion of each simply balanced the directional movement of attention through the acoustic spectrum across the experiment.

**Figure 1.**
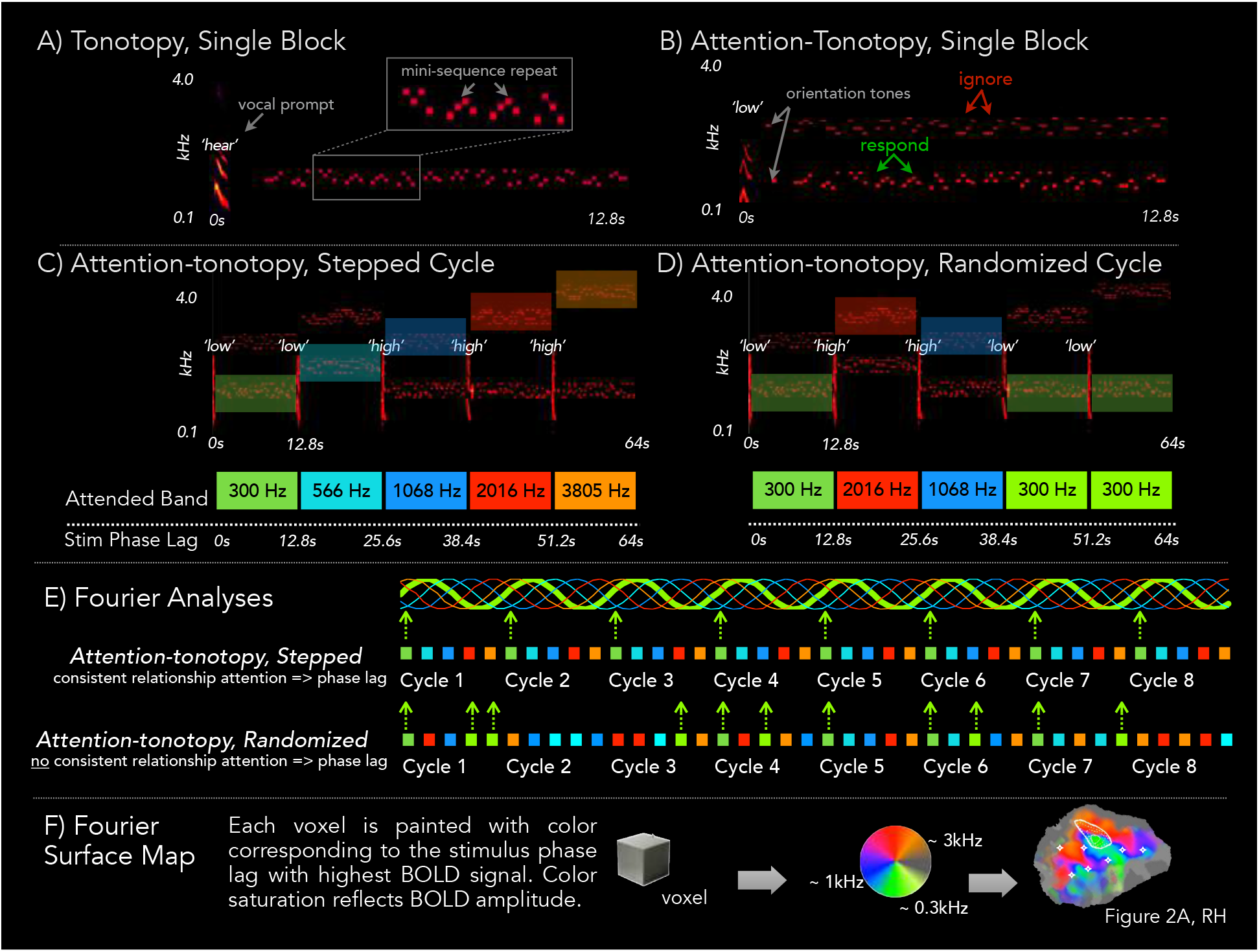
Stimuli and design overview. **(A)** In a representative 12.8s Tonotopy block a neutral verbal prompt *“hear”* precedes 14 four-tone mini-sequences sampled around one of 5 center frequencies. The task is to detect the 1-3 mini-sequence repeats embedded within the block. The grey box highlights a single mini-sequence repeat. **(B)** A single **Attention-Tonotopy (Attn-tono) block** includes two simultaneous streams of mini-sequences with distinct center frequencies. Mini-sequence repeats occur in each stream. A verbal prompt *(“high”* or *“low”*) directs listeners to attend to one stream and report mini-sequence repeats in that stream, while ignoring repeats in the unattended stream. Two randomly-ordered orientation tones at the center frequency of each stream alert listeners to the frequency neighborhood of the upcoming streams. **(C)** A single 64s cycle of Stepped Attn-tono blocks includes five 12.8s blocks that step up (shown), or down, in center frequency. In this single cycle, the frequency band to which attention is directed by the verbal prompt (indicated with *“high”/”low”* above each block) is acoustically matched with the Tonotopy cycle shown in **(B)**, but there are always competing unattended mini-sequences in a distinct frequency band. **(D)** A single 64s cycle of Randomized Attn-tono blocks is acoustically identical to the Stepped Attn-tono cycle in **(C)**, except that half of the verbal prompts have been swapped, and therefore no longer cue attention to frequency with consistent phase lags. **(E)** The distinction between Stepped and Randomized Attn-tono blocks is highlighted by examining the first three (of eight) **cycles of a Stepped (top) versus Randomized (bottom) Attn-tono run** The focus of attention is color coded in the frequency-band-specific manner shown in **(A)**. For the Stepped condition (top), there is a consistent relationship between the stimulus phase lag and the attended frequency across cycles within a run. Thus, for voxels that show a consistently higher response at one attended frequency band compared to all others, there will be a periodic response at 8 cycles/run at a given phase lag corresponding to the particular frequency band attended. For the Randomized condition (bottom), there is no consistent relationship, providing a control condition for Fourier analyses because frequency-band-directed attention is aperiodic across a run. **(F)** The stimulus phase lag with the highest periodic BOLD signal amplitude is determined for each voxel, mapped to the color wheel, and then painted onto the cortical surface patch. BOLD signal amplitude is mapped to the color’s saturation. (Note: In these panels A-D, stimulus intensity is adjusted across the spectrum to aid visual presentation of energy across frequency bands. See Methods for details on actual intensity across frequency bands).

In summary, in the *attn-tono* condition, attention alone was available to differentially drive responses to an approximately constant acoustic input, whereas in the *tonotopy* condition, responses were driven by spectrally-selective stimuli as well as by attention.

We analyzed mapping data using Fourier methods with individual and surface-based group analysis methods, as described previously (Sereno et al., 1995; Hagler et al., 2006; 2007). With this approach, voxels preferentially responding to a certain phase in a ‘stepped’ stimulus cycle are defined as those that have a significantly higher signal amplitude at this stimulus temporal frequency (meaning the slow frequency of the repeat of the spectral ramp) than at the average of other ‘noise’ frequencies (see Fig. 1e). Significant signal *phases* (a particular position in the cycle) are then mapped to a color wheel to indicate the voxel’s ‘best frequency’ and signal amplitude is mapped to the voxel’s color saturation (Fig. 1f). We time-reversed runs stepping down in frequency and averaged them with runs stepping up in frequency (Sereno et al., 1995; Talavage et al., 2004; Dick et al., 2012; Ahveninen et al., 2016). Cross-subject averaging of phase-encoded mapping data was performed using a method described previously (Hagler et al., 2007) in which the real and imaginary components of the signal with respect to the stepped cycle were sampled to the cortical surface and then averaged across subjects, preserving any phase information that was coherent over subjects.

Using previously established methods (Dick et al., 2012; Sereno et al., 2013; Lutti et al., 2014) see also (Glasser et al., 2016)), we used high-resolution quantitative multiparameter imaging to generate maps of estimated cortical myelination based on longitudinal relaxation times (quantitative T_1_). Recent work by multiple labs supports the hypothesis that T_1_ relaxation is reliably associated with quantitative differences in myelination in white matter and cortex (Sereno et al., 2013; Callaghan et al., 2014; Stüber et al., 2014; Dinse et al., 2015; Tardif et al., 2015; Turner, 2015; Tardif et al., 2016). Here, we calculated each subject’s R_1_ (1/T_1_) values, where the greater the R_1_, the higher the inferred myelin content. These R_1_ values were resampled onto his or her surface at a cortical depth fraction of 0.5, and also averaged across individuals using sulcus-aligned cortical-surface-based procedures (see below for further details).

### Participants

Eight adults (aged 23-45 y, mean 28 y; 6 female) participated; none reported a history of neurological disease or communication disorders. All had some childhood and/or adult musical training (one had a music degree), and had previous experience with longer scanning sessions. While musical training seemed to facilitate learning the experimental task, subsequent behavioral studies in the lab have shown that musically-naive subjects can also attain excellent performance with similar levels of training on this and even more demanding related tasks.

### Stimuli and Design

Stimuli were created using custom code in Matlab version 2015a (Mathworks, Inc.) and SoX version 14.4.2 (sourceforge.net). The basic stimulus unit was a four-tone *mini-sequence* (140 ms sine-wave tones including 10 ms linear amplitude ramp), with each tone drawn with replacement from a seven-semitone, band-delimited pool centered around one of five frequencies (300, 566, 1068, 2016, 3805 Hz; Fig. 1a). Fourteen mini-sequences formed a *block* (mean inter-sequence silent interval 240 ms, SD 10 ms). Each block contained one to three mini-sequence repeats (1:2:1 ratio of 1, 2, and 3 repeats). When there was more than one repeat per block, mini-sequence repeat pairs were separated by at least one intervening mini-sequence. Each block began with a verbal prompt: *‘hear’, ‘high’*, or *‘low’;* loudness-equalized Mac ‘Victoria’ voice, mean duration 506 ms (SD 36 ms), padded with silence to 800 ms total duration. This prompt was followed by 800 ms silent gap (tonotopy) or tone-cue (attn-tono), then the 14 mini-sequences (11.2s in total), for a total block duration of 12.8s.

The task was to detect mini-sequence repeats in the attended frequency band (i.e., a 1-back task). In the tonotopy condition, mini-sequences were confined to a single frequency band preceded by the neutral verbal prompt *‘hear’* (Fig. 1a). In two of the four single-band tonotopy runs, block center frequency was stepped from low-to-high over a 64-second cycle with 8 cycles/run; step direction was reversed (high-to-low) for the other two runs. This is a ‘discrete’ version of phase-encoded designs commonly used in visual, somatosensory, and auditory mapping studies (Engel et al., 1994; Sereno et al., 1995; Da Costa et al., 2011; Dick et al., 2012; Langers and van Dijk, 2012; Langers et al., 2014; Saenz and Langers, 2014).

The attn-tono condition had the exact mini-sequence patterns from the tonotopy blocks, but there also were simultaneous, competing mini-sequences in a distinct frequency band with a center frequency at least 14 semitones apart (Fig. 1b; 300 vs. 1068 Hz; 300 vs. 2016 Hz; 300 vs. 3805 Hz; 566 vs. 2016 Hz; 566 vs. 3805 Hz; 1068 vs. 3805 Hz; not all center frequencies were paired due to the 14-semitone constraint). The verbal prompt (*high* or *low*) initiating each attn-tono block signaled participants to perform the 1-back task on either the higher or lower frequency band. Immediately after the verbal prompt, a randomly-ordered pair of sine-wave tones cued the center frequencies of the upcoming block (140 ms tones including 8 ms linear on/off amplitude ramp; 26 ms inter-tone silence, tone pair followed by 494 ms silence, total duration 800 ms). Crucially, there were mini-sequence repeats even in the unattended band to assure that attention directed to the task was endogenously driven rather than being attracted by stimulus repetition effects (Barascud et al., 2016).

There were two attn-tono conditions: ‘stepped’ and ‘randomized’. Analogous to single-band tonotopy runs, in stepped attn-tono runs the verbally-cued frequency band implicitly stepped up (2 runs) or down (2 runs) in frequency over a 64-sec *cycle* (Fig. 1c). This cued iterative stepping through the frequency spectrum facilitates transfer of attention to each frequency band (as in traditional phase-encoded designs) and supports Fourier approaches to analysis (Fig. 1e). Each randomized attn-tono run was acoustically identical to a stepped run, but the verbal prompt was manipulated so that there was no systematic, stepped organization of mini-sequence center frequencies through the spectrum (Fig. 1d). For this condition, frequency bands were cued at inconsistent phase lags within the 8 cycles/run, thereby phase-canceling any periodic attentional response; this is schematized in Fig 1e. This randomized-order control is important, as there is a small (~1 octave) overall shift in spectral mean over the course of an attn-tono stimulus cycle that is unavoidable due to the constraints on the pairing of frequency bands.

Each of the twelve 9.6-minute-long runs was composed of eight 64s cycles plus 32-sec silent periods at the beginning and end of each run to allow for calculation of baseline auditory activation (Klein et al., 2014).

### Behavioral Thresholds and Training

Participants first underwent behavioral tests of monaural pure-tone thresholds and binaural thresholds for detecting mini-sequences in quiet and in acoustic noise generated by the MRI scanner running the multiband EPI sequence. This provided a basis for adjusting center frequency amplitudes to approximate equal loudness in scanner noise. Participants also trained on the mini-sequence detection task in quiet and in acoustic scanner noise across two sessions.

### Imaging Data Acquisition

Structural and functional images were acquired on a 3-Tesla Siemens Verio wide-bore MRI scanner at the Scientific Imaging and Brain Research (SIBR) Center at Carnegie Mellon University using a phased array 32-channel head coil across three scan sessions on separate days. Stimulus presentation was under the control of a MacPro running PsychToolbox 3.0.12 in Matlab (The Mathworks, Inc.), with audio output to an external AD/DA converter (Babyface, RME) connected to an amplifier (Pylepro) that delivered stimuli to participants in the scanner diotically over MRI-compatible earbuds (Sensimetrics S14). All stimuli were pre-filtered to equalize sound stimuli according to the earbuds’ frequency response. After participants were settled into the bore, sound volume was adjusted so that participants could comfortably hear all frequencies through scanner noise. Participants wore a fiber optic response glove (Current Designs) that communicated with a Brain Logics I/O device (Psychology Software Tools, Inc); participants used the glove to respond to mini-sequence repeats using the right index finger. During all functional scans, subjects closed their eyes to reduce the potential for stimulus-correlated eye movements.

In the initial scanning session (~50 min), we acquired multi-parameter mapping (MPM) images for quantitative myelin mapping and structural identification of primary auditory cortex on an individual basis while participants watched a film. Proton density-weighted (PDw), T1-weighted (T1w), and magnetization transfer (MTw) images were acquired using an in-house 3D FLASH pulse sequence (voxel size: 0.8 × 0.8 × 0.8 mm^3^, matrix = 320 × 280 × 208, TR = 25.0 ms, bandwidth 488 Hz/px, excitation flip angle: 6° (PDw/MTw) or 21° (T1w), slab rotation 30°). To accelerate this high resolution acquisition, a partial Fourier acquisition (6/8 coverage) was used in the inner phase-encoded direction (RL) and parallel imaging was used along the outer phase encoding direction (AP), reconstructed using the GRAPPA algorithm (acceleration factor 2, 18 integrated auto-calibration lines) as implemented on the scanner platform. Four gradient echoes were acquired for each contrast (TE=2.5, 4.74, 6.98, 9.22 ms) after each excitation pulse and averaged to improve SNR (Helms et al., 2009). Each FLASH acquisition lasted 9 minutes 45 seconds. Quantitative R_1_ (=1/T1) maps were estimated from the PDw and T1w images according to the model developed by Helms et al. (Helms et al., 2008) including a correction for RF transmit field inhomogeneities (Lutti et al., 2010) and imperfect spoiling (Preibisch and Deichmann, 2009). The transmit field map was calculated using a 3D EPI spin-echo (SE)/ stimulated echo (STE) method (Lutti et al., 2010; 2012); FOV = 256 × 192 × 192 mm, matrix = 64 × 64 × 48, TE = 53.14 ms, TM = 47.60 ms, TR = 500 ms, bandwidth = 2298, nominal α varying from 135° to 65° in steps of 5°, acquisition time 6 minutes) and was corrected for off-resonance effects using a standard B0 field map (double gradient echo FLASH, 3x3x2 mm isotropic resolution, whole-brain coverage).

The final two scanning sessions acquired functional data for four runs each of the tonotopy, ‘stepped’ attn-tono, and ‘randomized’ attn-tono conditions. The runs were interleaved across conditions and designed to assess phase-encoded functional influences of selective attention across frequency (‘stepped’ attn-tono), the functional response to identical acoustics without systematic phase encoded shifts of attention (‘randomized’ attn-tono), and functional responses to single frequency bands identical to the attended bands in attn-tono, with phase-encoded steps through frequency and no distractor frequency bands (tonotopy). Across all functional runs, participants engaged in detecting repeats (1-back) of the four-tone mini-sequences. Run order was counterbalanced according to condition and whether the cycle involved steps up or down in frequency.

Functional images were acquired using a T2*-weighted echo-planar imaging (EPI) pulse sequence (44 oblique axial slices, in plane resolution 3 mm × 3 mm, 3 mm slice thickness, no gap, repetition time TR = 1000 ms, echo time TE = 41 ms, flip angle = 61°, matrix size = 64 × 64, field of view FOV = 192 mm). All EPI functional scans were performed using 4x multi-band acceleration (Feinberg et al., 2010; Feinberg and Setsompop, 2013). There were 584 repetitions acquired per run, with the first 8 images discarded to allow for longitudinal magnetization to arrive at equilibrium. Runs were pseudo-randomly ordered across participants.

### Image Preprocessing

#### Cortical surface creation, and mapping of R_1_values

Each subject’s cortical surface was reconstructed from a contrast-optimized synthetic FLASH volume, created with mri_synthesize in Freesurfer from scaled and truncated versions of the T1 and proton-density volumes; another MPRAGE-weighted version was created for use with the automated Freesurfer Talairach procedure. Both volumes were conformed to 1mm isotropic resolution and used in a customized reconstruction pipeline version. (In particular, the subject’s PD volume was used to deskull the synthetic FLASH image using a ‘shrink-wrap’ technique (Dale and Sereno, 1993)). After inspection for reconstruction quality, R_1_ values were resampled from 50% cortical depth fraction to the subject’s surface, and also morphed to the unit icosahedron for cross-subject curvature-aligned cortical surface based averaging (Fischl et al., 1999).

#### EPI processing

Each functional image from both sessions was aligned to a reference volume from the middle of the first run using AFNI’s 3dvolreg; registration and motion correction goodness were hand-checked for each run. The reference volume was aligned to the subject’s cortical surface using boundary-based registration in Freesurfer (Greve and Fischl, 2009), verified using manual blink comparison, and applied to the volume-aligned EPI data for resampling. EPI data were analyzed in native space without any spatial smoothing using both Fourier and general linear methods.

### Experimental Design and Statistical Analysis

As noted above, the fMRI experiment used a ‘discrete’ version of a traditional phase-encoded design, such that both Fourier-based and general linear model approaches could be used. Fourier analyses were carried out in csurf (http://www.cogsci.ucsd.edu/~sereno/.tmp/dist/csurf) with individual and group analysis methods employed as previously described (Sereno et al., 1995; Sereno and Huang, 2006; Hagler et al., 2007). Functional activation amplitude was estimated as the Fourier amplitude of the periodic BOLD signal (proportional to percent response) at the frequency of the stimulus cycle (8 repetitions per run). An F statistic was calculated by comparing that amplitude to the average amplitude of other ‘noise’ frequencies (Hagler et al., 2007). Periodic signal components with very low frequencies (due to slow head motion) and the second and third harmonic of the stimulus were excluded as neither signal nor noise (this is mathematically equivalent to first linearly regressing out these frequencies as nuisance variables before calculating significance). The phase of the signal, which corresponds to a particular point of the stimulus cycle, was then mapped to the color wheel and the amplitude of the signal at each vertex was mapped to color saturation (Gouraud sharing within each face). Runs with downward frequency steps were time-reversed and averaged with upward-stepped scans in order to cancel fixed voxel-specific delays in the BOLD response.

Linear modeling was carried out in FSL (Smith et al., 2004). For all runs, the motion-registered data were high-pass-filtered (100 sec) and prewhitened; a hemodynamic model corresponding to each stimulated and attended (tonotopy condition) or attended (stepped, randomized attn-tono conditions) frequency band was created by convolving the 12.8-sec block with a gamma function (lag 6s, SD sc). In a separate multiple regression, the unattended (ignored) frequency band was modeled for both stepped and randomized attn-tono conditions. The verbal cue was also modeled; all models were temporally filtered before multiple regression. Coefficients from the first-level contrasts for each of the four runs were combined in a fixed-effects analysis for each condition; data from the stepped and random block conditions were also combined in an eight-run average.

Cross-subject averaging of phase-encoded mapping data was performed using the methodology developed by Hagler and Sereno (Hagler and Sereno, 2006) in which the real and imaginary components of the signal with respect to the stimulus ramp are averaged across subjects, preserving any phase information consistent between subjects. This was performed by projecting each participant’s phase-encoded map to the FreeSurfer spherical atlas using mri_surf2surf, performing 1 step of surface-based smoothing (< 1 mm FWHM in 2D), averaging across subjects at each vertex, then painting back onto a single subject’s surface for viewing.

For the multiple regression analyses, the same sampling process was used to sample each subject’s contrast parameter estimates for cross-subject averaging and t-tests.

Surface-based cluster exclusion was used to correct for multiple comparisons in the groupwise averages (surfclust and randsurfclust from (Hagler et al., 2006)). The exclusion criterion (only surface clusters > 78 mm^2^ unless otherwise noted) was determined based on the minimum estimated cortical area from iterative random sampling of cluster sizes (N=10000 iterations per hemisphere in randsurfclust) required to achieve a corrected alpha of p < 0.001 for each hemisphere, based on an initial uncorrected alpha of vertexwise p < 0.01.

#### ROI analyses

We quantified the similarity between frequency band response profiles driven by stimulus+attention (tonotopy) versus attention alone (attn-tono) in a ‘quilt’ of small cortical-surface-based ROIs that tiled the temporal plane. ROIs (as seen in Fig. 4c) were created by on a single subject’s right and left hemisphere flattened patches by flooding all vertices within a 4mm radius around a central selected vertex. Each of the ROIs (57 in the right hemisphere patch, and 68 in the slightly larger left patch) were then spherically morphed to the other 7 subjects’ flattened patches. Spurious ROI sampling on the edges of the patches was manually corrected on the original subject’s inflated surface and re-morphed to all other subjects. Each ROI was then projected into the registered native-space EPI volume using Freesurfer’s mri_label2vol (sampled from the grey-white boundary to 0.8 of the calculated cortical depth, with fillthresh set to 0.5). For each subject, within each ROI, we calculated the average parameter estimate for each frequency band for tonotopy, and combined stepped and randomized attn-tono conditions. For each ROI, we then ran a linear model with average tonotopy parameter estimates for the 5 frequency bands predicting average attn-tono parameter estimates for the same bands, including subjects as a random factor. The resulting partial t-statistic for each ROI was z-transformed and color-rendered in Fig. 4c, with p-value thresholds Bonferroni-corrected to p < 0.05 for the number of ROIs per hemisphere, and indicated by the white outline surrounding the set of ROIs that surpass this z-threshold.

## Results

### Fourier-based Analyses

#### Stimulus - and attentionally-driven tonotopic organization in human auditory core

As a necessary first step, we characterized basic tonotopic (stimulus-driven) organization in and immediately around myelin-defined auditory core. The group-average R_1_-based estimates of myelination (inflated hemispheres in left-most panel of Fig. 2) show that the highest R_1_ values occur within primary somatomotor areas along the central sulcus, and in the typically keyhole-shaped ‘auditory core’ lying along and immediately surrounding Heschl’s gyrus. Its shape is more easily observed in the auditory cortex flat patches shown in Figure 2a-c, depicted with isointensity R_1_ contours, with Heschl’s gyrus marked for reference. The group-averaged topography of preferred frequency around auditory core has a typical arrangement (Dick et al., 2012; De Martino et al., 2015b), with the core surrounded by a high-frequency ‘V’. Preferred frequency descends into the center of core (where R_1_ values are highest) before reversing and slowly ascending to mid-frequency preferred frequencies anterolaterally (and to some extent posterolaterally). Fig. 3 shows tonotopic maps for each individual listener. In general, the relationship between auditory core and tonotopy group is conserved across listeners, albeit with some variability in the shape and extent of the isointensity R_1_ contours.

**Figure 2.**
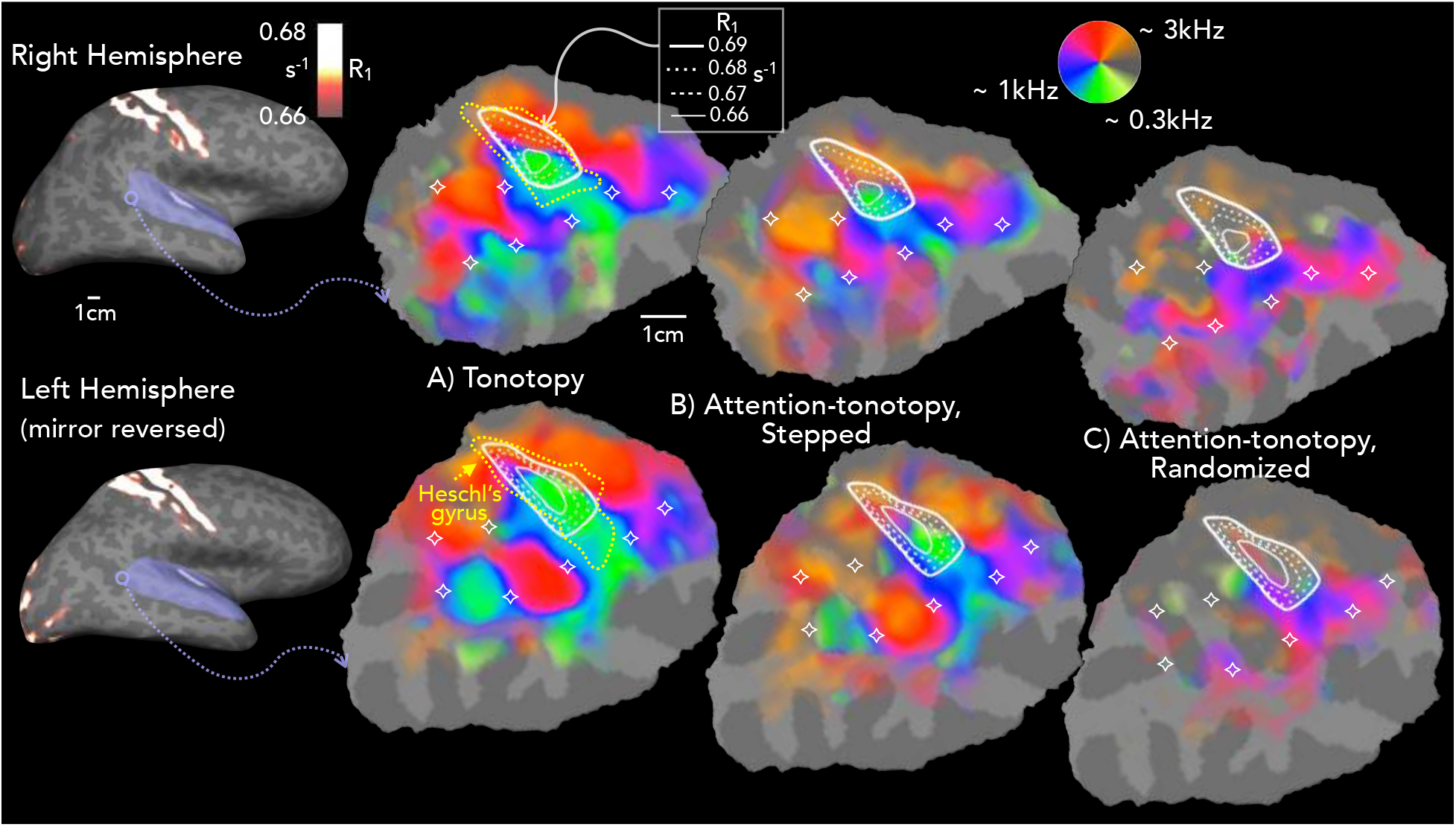
Group activation for Tonotopy and Attention-Tonotopy conditions, with R_1_ contours showing putative auditory core. The left-most panel shows cortical-surface-based group-averaged R_1_, projected on the lateral inflated surface of one subject. (The left hemisphere is mirror-reversed to align cortical maps for visual comparison). For tonotopic map display, a patch of cortex including the entire temporal plane (shown in purple on the inflated surface) was cut and flattened. Panels A-C show this region enlarged, with isocontour lines showing quantitative R_1_ values for the group-averaged putative auditory core, and color maps showing group-averaged best frequency as a function of (**A**) Tonotopy, (**B**) Attn-tono (Stepped), and (**C**) Attn-tono (Randomized) conditions. The stars are fiduciary points to assist in visual comparisons of maps across conditions; the outline of Heschl’s gyrus is in yellow dashed lines (in (**A**), from the individual subject whose cortical patch was used). Consistent with previous work, the tonotopic map is characterized by two pairs of three interlacing best - frequency ‘fingers,’ with the high-frequency fingers (red/orange colormap) showing greatest frequency preference medially and extending laterally, where they meet interdigitated lower-frequency fingers (green/yellow colormap) extending lateral to medial, with the longest ‘middle’ lower-frequency finger extending about halfway into auditory core. This pattern is evident in Fourier-analysis-derived maps of the ‘stepped’ Attn-tono condition but not in the ‘randomized’ Attn-tono condition, for which the attentional response was phase-cancelled. All maps are statistically masked by overall activation to tonotopy in each hemisphere (cluster-corrected p < 10^−8^, and gently shaded to show relative amplitude).

We then asked whether ‘attention-tonotopic’ mapping resembled the tonotopic case in and around auditory core. Here, the group-level spatial distribution of tonotopy is closely recapitulated when spectrally-directed attention (‘stepped’ attn-tono condition) alone modulates activation (Fig. 2b). This holds true in and around the keyhole-shaped hyperintensity defining core, with a slight exception in the transition from higher to lower frequency preference in mid core. In contrast, the group-level ‘randomized’ attn-tono response is much weaker, with almost no correspondence with the tonotopic map (Fig. 2c), despite being acoustically identical to ‘stepped’ attention-o-tonotopy but for the shuffling of the verbal prompt ordering that destroyed the consistent phase-lag associated with specific frequency bands supporting these Fourier-based analyses. The one potential exception is in and around posterolateral core, where there is a low-to-mid frequency progression that is similar in attention-o-tonotopic and tonotopic maps, particularly in the left hemisphere. (This may be due to the small (~1 octave) overall shift in spectral mean over the course of a stimulus cycle noted in Methods).

#### Stimulus-and attentionally-driven tonotopic organization outside of auditory core

In line with results from previous fMRI studies (Talavage et al., 2004; Woods et al., 2009; Humphries et al., 2010; Barton et al., 2012; Dick et al., 2012; Moerel et al., 2012; Saenz and Langers, 2014; Thomas et al., 2015; De Martino et al., 2015b; Ahveninen et al., 2016; Leaver and Rauschecker, 2016; Riecke et al., 2016), there is stimulus-driven tonotopic mapping extending well beyond auditory core, spanning the temporal plane and continuing into the superior temporal sulcus (STS). As shown in Fig. 2a, the overall arrangement is characterized by two pairs of three interlacing best-frequency ‘fingers,’ with the high-frequency fingers (red/orange colormap) predominating medially and extending laterally, where they meet interdigitated lower-frequency fingers (green/yellow colormap) extending lateral to medial, with the longest ‘middle’ lower-frequency finger extending about halfway into auditory core. Similar to tonotopy within auditory core, the overall pattern of group activation can be observed in the majority of individual subjects (Fig. 3), but there is also considerable individual variability in the complexity, topography, and extent of tonotopic and attn-tono mapping, similar to that observed in the fMRI studies cited above (as well as electrophysiological studies in a number of studies in macaque and owl monkey, e.g., (Merzenich and Brugge, 1973; Morel et al., 1993)).

As can be seen in the maps in Fig. 2b, the tonotopically-aligned maps evoked by spectrally-directed attention are also present in the majority of auditory cortex outside of auditory core. Again, the structure of the tonotopic map (as revealed by Fourier analysis) is abolished when the attentional cue is randomized, thereby eliminating any consistent relationship between attended frequency band and phase lag.

The similarity between the maps evoked by presentation of a single frequency band (tonotopy) versus attention to one of two simultaneously-presented frequency bands (‘stepped’ attn-tono) can also be seen in each individual subject (Fig. 3). As with the group-averaged data, there is a close correspondence in the progression of preferred frequencies across auditory cortex in individual subjects. The similarity between the tonotopic and attn-tono maps is particularly striking in subjects 1, 2, 5, 6, and 7. The tonotopic organization of individual subjects demonstrated overall commonalities, but with notable differences, even between individual subjects’ right and left hemispheres, particularly outside of auditory core (Humphries et al., 2010; Moerel et al., 2014; Saenz and Langers, 2014). However, individual peculiarities were replicated across tonotopic and attn-tono conditions. In some subjects, there was a surprising lack of strong tonotopic mapping (subject 3 for which poor tonotopy may be due to greater EPI warping, and also subject 4 for which low frequencies dominated the tonotopic maps). In summary, there was a strong correspondence between tonotopic and attn-tono maps at both the group and individual levels.

**Figure 3.**
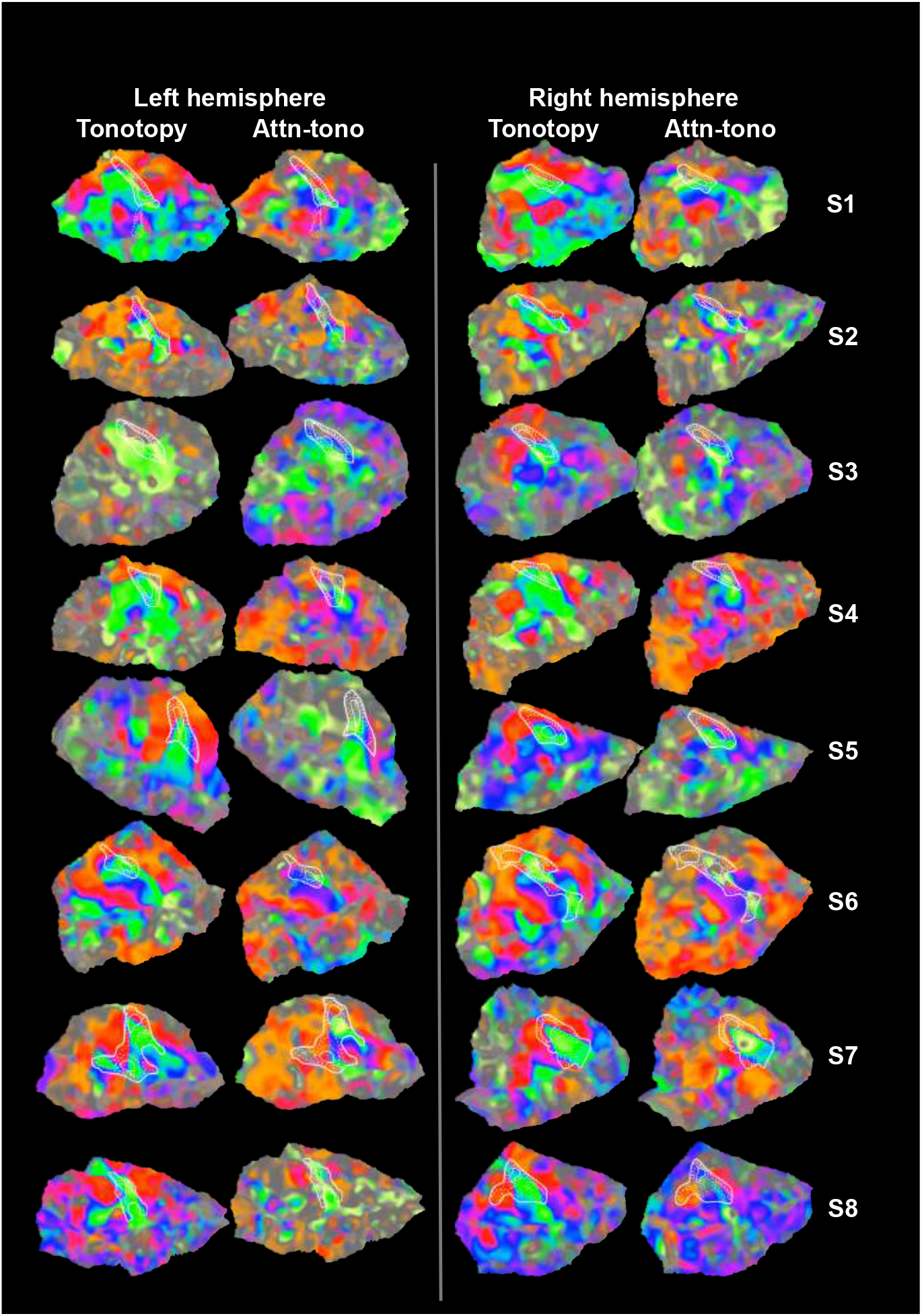
Individual subjects’ Tonotopy and Attention-Tonotopy maps. Each subject’s Tonotopic and Attn-Tono (Stepped) Fourier-analysis-derived maps are displayed on the same subject’s flattened superior temporal cortical patch. R_1_ isocontours around presumptive auditory core are shown in white, with thick solid lines depicting the lowest-valued (outermost) R_1_ isocontour, thin solid lines depicting the highest (innermost) R_1_ isocontour, and dashed lines showing intermediate values. (R_1_ values differ somewhat across individuals). Activation maps are gently shaded to show changes in response amplitude, but are unthresholded for comparison with individual maps from previous studies, e.g., Da Costa et al., (2011).

### Multiple regression analyses

#### Winner-Takes-All: Maps of ‘stepped’ versus ‘randomized’ attention conditions, and quantitative concordance of tonotopic and attn-tono maps

In a complementary analysis, we used standard multiple regression techniques (see Methods) to estimate the BOLD response to each center-frequency band when it was presented in isolation (tonotopy), versus when it was attended in the presence of a distractor band (attn-tono). This allowed us to make use of the attentionally-driven signal in the randomized attn-tono condition and to combine these data with the results from the stepped attn-tono condition to increase statistical power. It also allowed us to verify that the attention effects generalize when listeners direct attention without the ‘crutch’ of consistent stepping up or down across attended frequency bands.

**Figure 4.**
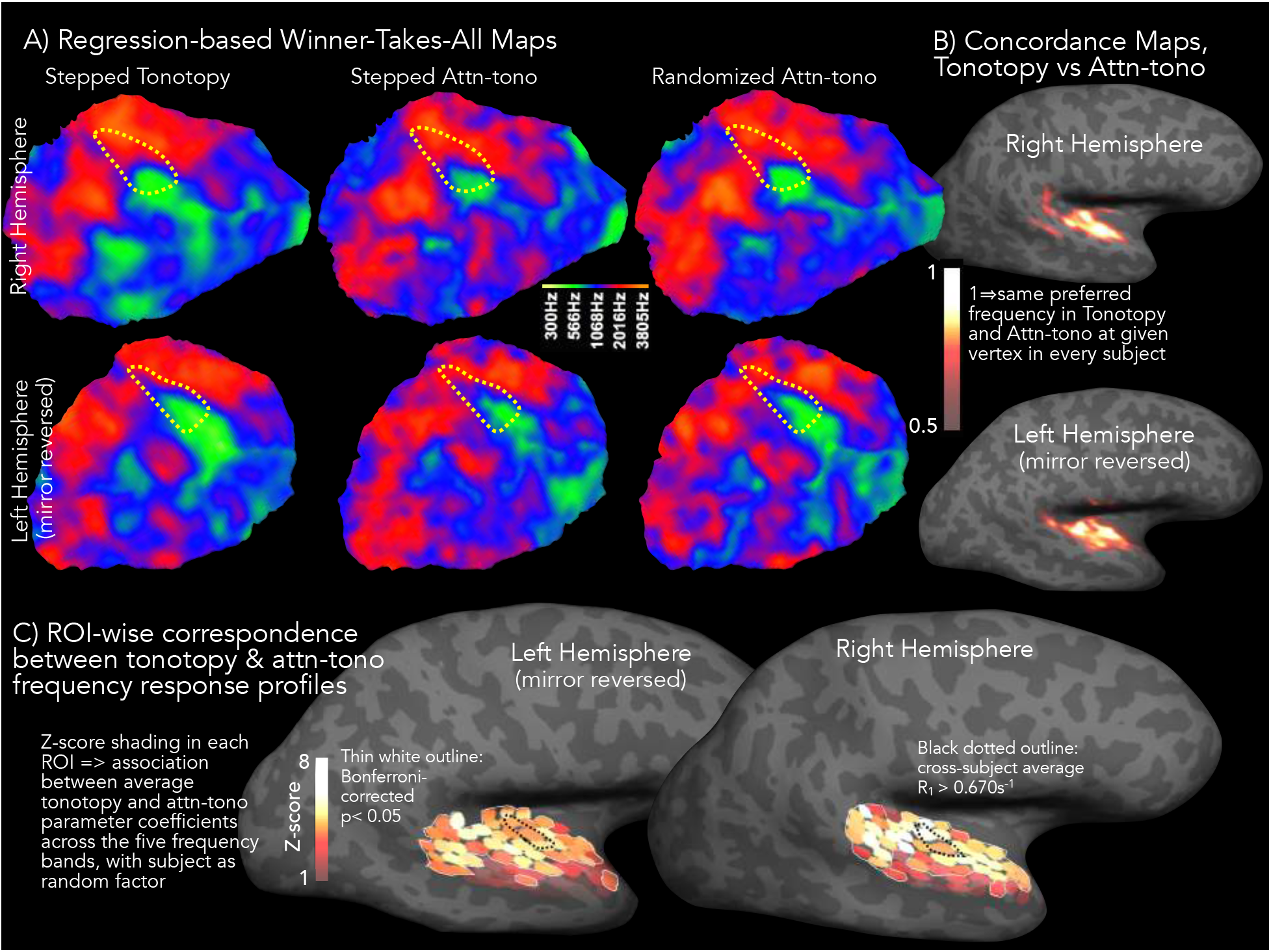
Comparison of responses in regression-based ‘Winner-Takes-All’ maps, Tonotopy and Attn-Tono. (A) The color maps projected onto the right (*top panels*) and left (*bottom panels*) hemisphere cortical patches (same as patches shown in purple in Figure 2) show the cross-subject average best frequency band (‘Winner-Takes-All,’ or WTA) for ‘stepped’ tonotopic (left) ‘stepped’ attn-tono (middle) and ‘randomized’ attn-tono conditions. The dotted yellow line depicts the outermost R_1_ contour (0.68 sec^-1^) around presumptive auditory core as shown in Figure 2. **(B)** ‘Concordance maps’ are rendered in heatscale on the inflated hemispheres to illustrate the similarity in best frequency between tonotopic and attn-tono maps (the latter averaged over stepped and randomized blocks). These maps were calculated in two stages. First, in each subject’s native EPI space, a voxel was coded as ‘1’ if tonotopic and attn-tono stimuli evoked the same best frequency, and otherwise coded as ‘0’. Second, for each subject, the concordance maps were resampled to the individual’s cortical surface, and projected onto the unit icosahedron for cross-subject surface-based averaging, thereby creating a composite measure of agreement between tonotopic and attn-tono maps, weighted by the consistency of this agreement across subjects. The concordance maps are statistically masked with a cross-subject t-map, calculated versus chance agreement (p=0.20) with a surface cluster correction of p < 0.001 (vertex-wise p<0.05, cluster threshold surface area> 203 mm^2^, (Hagler et al., 2006)). **(C)** The shading in each small ROI patch shows the z-score for the partial fit between tonotopy and attn-tono responses to each frequency band (with subjects as a random factor). ROIs with significant z-scores (Bonferroni-corrected p-value threshold of p < 0.05) are indicated by the thin white outline.

The auditory cortical patches in Fig. 4a show the cross-subject average ‘Winner-Takes-All’ (WTA) best frequency band (most positive-going BOLD response relative to resting baseline) maps for tonotopy and attn-tono conditions (with no shading for response amplitude). These are overlaid with the outermost R_1_ isocontour (dashed yellow) corresponding to auditory core. As should be expected, the topography of the WTA maps essentially recapitulates the topography revealed by the phase-encoded analyses. The same holds true of the attn-tono WTA maps from both the ‘stepped’ and importantly, the ‘randomized’ block conditions (Fig. 4a); this result confirms that even without the ‘crutch’ of the stepping frequency band, listeners can direct their attention to specific frequency bands.

The WTA approach also allowed us to straightforwardly quantify the within-subject correspondence between voxel-wise best frequency, as estimated by tonotopy and by attn-tono. Here, we coded each voxel in native space as a ‘1’ when best frequency was identical in both conditions, and a ‘0’ otherwise. We then resampled each subject’s binary maps to their cortical surface, and then averaged across subjects to create a ‘concordance’ map (Fig. 3b). These maps (statistically thresholded at vertex-wise p< 0.01, with surface-cluster-corrected alpha of p < 0.001) show that across subjects there was high concordance across best frequency maps evoked by stimulus and by attention across much of the temporal plane in both hemispheres, with little concordance in non-auditory areas.

### Comparison of response profiles to all frequency bands across tonotopy and attn-tono

As has been shown previously, e.g., Moerel et al., (2013), hemodynamic responses to frequency in auditory cortex are not necessarily bandpass, but can more complex and multipeaked. Therefore, we also examined whether attention to a given frequency band in the presence of a distractor band recapitulates the more graded response to non-preferred frequencies observed when that frequency band is presented in isolation. To do this, we created and surface-morphed a set of small cortical ROIs to each subject (see Fig. 4c and Methods), and quantified the similarity between the tonotopy and attn-tono response profiles in each ROI in each hemisphere by regressing the mean tonotopic parameter estimate for each frequency band against the attn-tono parameter estimate (with subjects as a random factor).

Converging with the results from the ‘concordance’ maps from the Winner-Takes-All analyses (Fig. 4b), the ROI analyses (Fig. 4c) show that individual subjects’ tonotopy and attn-tono responses profiles are significantly associated across most of auditory cortex (all ROIs within the white border), with the exception of the most lateral aspects of the STG and upper bank of the STS. (Note that while there is a strong relationship between tonotopy and attn-tono response profiles of each subject within a given ROI, there is cross-subject variability in the particular shape of those response profiles, as suggested by the individual maps in Fig. 3). There is a broad tendency for tonotopy/attn-tono profile similarity to be strongest posterior-medially in both hemispheres, and no clear indication that profile similarity is higher in auditory core (indeed, this is not the case in the left hemisphere).

### Loser-Takes-All: Maps of ‘dis-preferred’ frequency

Given the graded nature of frequency response preferences we observed, we suspected that there would be a large-scale topography associated with the minimum BOLD response across frequency, and that this topography would also be recapitulated by attention. Thus, we also performed a parallel ‘Loser-Takes-All’ (LTA) analysis, in which we coded voxels by the frequency band driving the minimum BOLD response (again relative to resting baseline) and analyzed as above. The average descriptive LTA maps show roughly opposite frequency responses compared to the WTA tonotopic maps, with higher-frequency-band-preferring regions in the tonotopic map being least driven by lower-frequency-bands, and vice versa (Fig. 5a). (There is also some overlap in the ‘mid-frequency-preferring’ regions, likely due to blurring of values when averaging subjects’ integer-based maps). There is also quite close correspondence between the frequency band evoking the least response in the tonotopy (stimulus) condition and the smallest BOLD response evoked by attending to a given frequency band. The LTA concordance maps show that in the right hemisphere, the alignment of tonotopic and attn-tono maps is greatest in more lateral and anterior auditory cortex, with qualitatively somewhat greater concordance more medially in the left hemisphere (Fig. 5b, statistical thresholding as in Fig. 4b).

**Figure 5.**
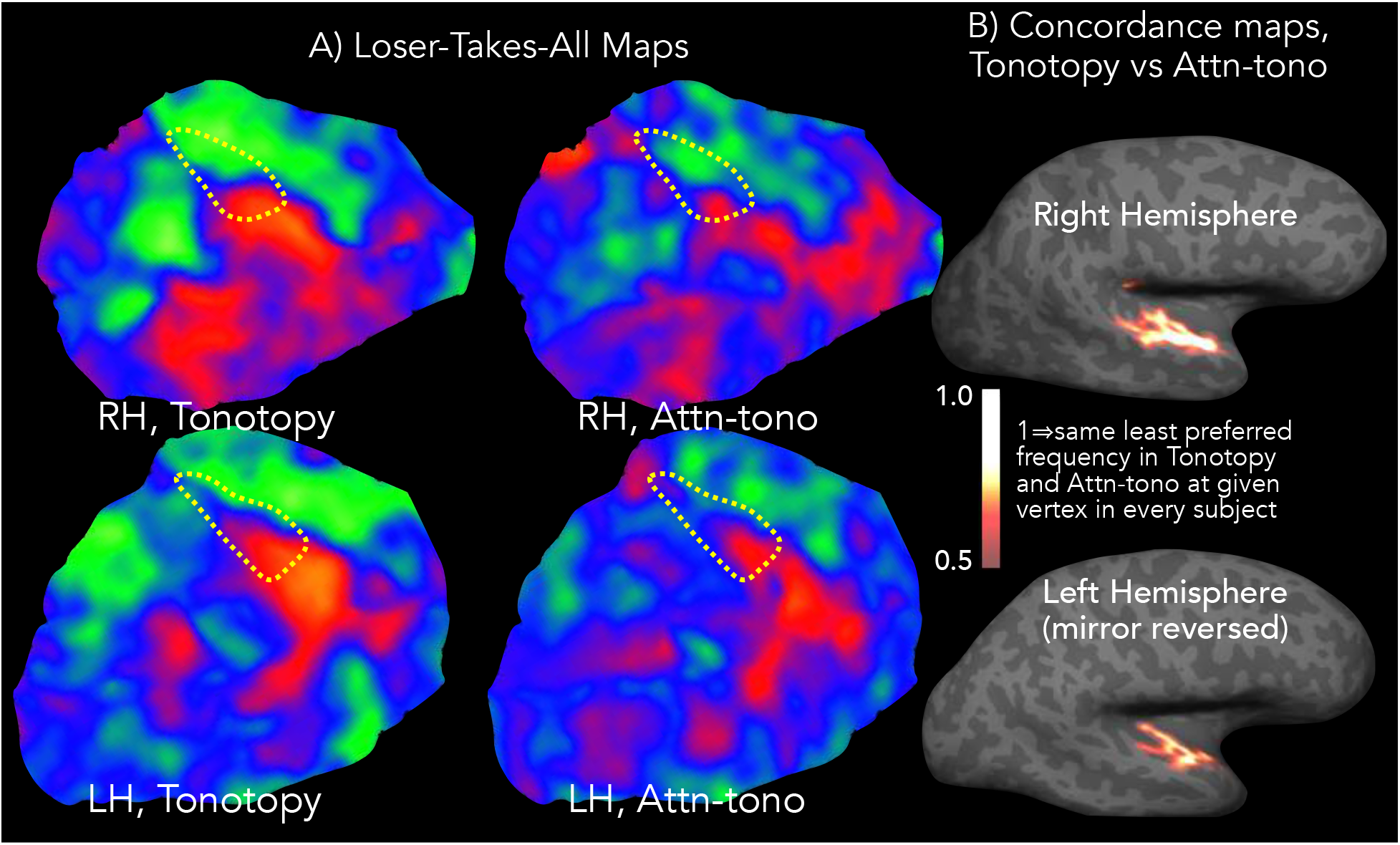
Comparison of responses in regression-based ‘Loser-Takes-All’ maps, Tonotopy and Attn-tono. (**A**) The colormaps projected onto the same cortical patches as Figures 2 and 4 show cross-subject group-average maps that depict the frequency band that drives the *least activation* compared to all other frequency bands (‘Loser-Takes-All’, or LTA) in tonotopy and attn-tono (stepped plus randomized blocks) conditions and in right and left hemispheres. As in Figure 4, the presumptive auditory core shown by the dashed yellow line depicting the outermost R_1_ contour (0.68 sec^-1^). **(B)** The tonotopic versus attn-tono LTA concordance map was created as in Figure 4b; note that the midpoint of the heatscale has been lowered slightly compared to Figure 4b. The dotted yellow R_1_ isocontour is the same as Figure 4a.

### Difference in activation across auditory areas when best frequency is attended versus ignored

We also assessed the strength and consistency of BOLD-related frequency-band-selective attention across subjects. We first used a subject’s native-space WTA map to establish each voxel’s best frequency. Then, we assigned each voxel the parameter estimate for the difference in activation between attending to its best frequency in the presence of a distractor, versus attending to the distractor and ignoring its best frequency. (In other words, the value at each voxel was the estimated difference in activation between attending to, versus ignoring, its best frequency in the presence of other frequency bands). We then resampled each subject’s native-space ‘attention map’ to her/his cortical surface to allow for surface-based cross-subject averaging and statistical testing (again with a vertex-wise p< 0.01 threshold and surface-cluster-corrected alpha of p < 0.001). Fig. 6 shows that across subjects there was significantly greater activation across most of auditory cortex when best frequency was attended versus ignored. The widespread attention effect included all of R_1_-estimated auditory core (outlined in green), extending from the inferior circular sulcus laterally to the upper bank of the STS, and antero-posteriorly from the temporal pole onto the planum temporale.

**Figure 6.**
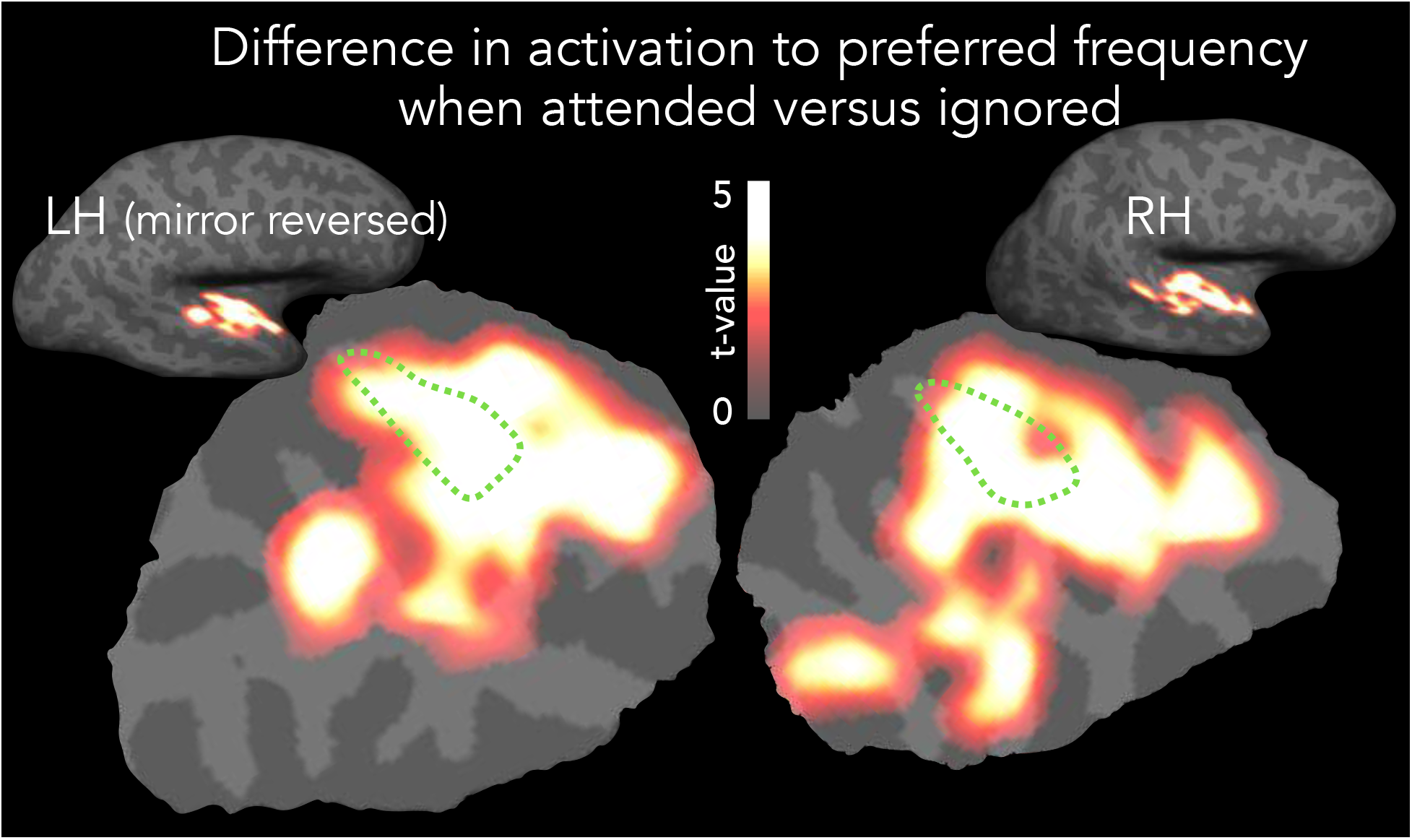
Comparison of maps when best frequency is attended versus ignored. The heatscale (t-values, thresholded as in Figure 4b) depicts the cross-subject cortical-surface-based average difference in activation when the subject-specific best frequency band of each voxel was attended versus ignored. The dotted green R_1_ isocontour estimating auditory core is as in Figure 4a.

### Relationship of tonotopic and attn-tono map strength to MR-estimated myeloarchitecture

Typically, assays of cortical myelination are used to differentiate the most highly-myelinated cortical regions (like auditory core, MT/V5, or V1) from adjacent regions. This is true whether cortical myelination is assessed using ex-vivo ‘gold-standard’ approaches such as Gallyas staining, or estimated through *in-vivo* MRI T1/T2 ratio, quantitative R_1_, or magnetization transfer measures. However, more subtle myelination changes that occur throughout cortex may spatially correspond with changes in functional characteristics (Glasser et al., 2016; Wallace et al., 2016). For instance, recent combined fMRI and high-resolution quantitative MR show that slight reductions in cortical myelination in primary somatomotor cortex reliably occur at the border between face and hand areas (Kuehn et al., 2017).

**Figure 7.**
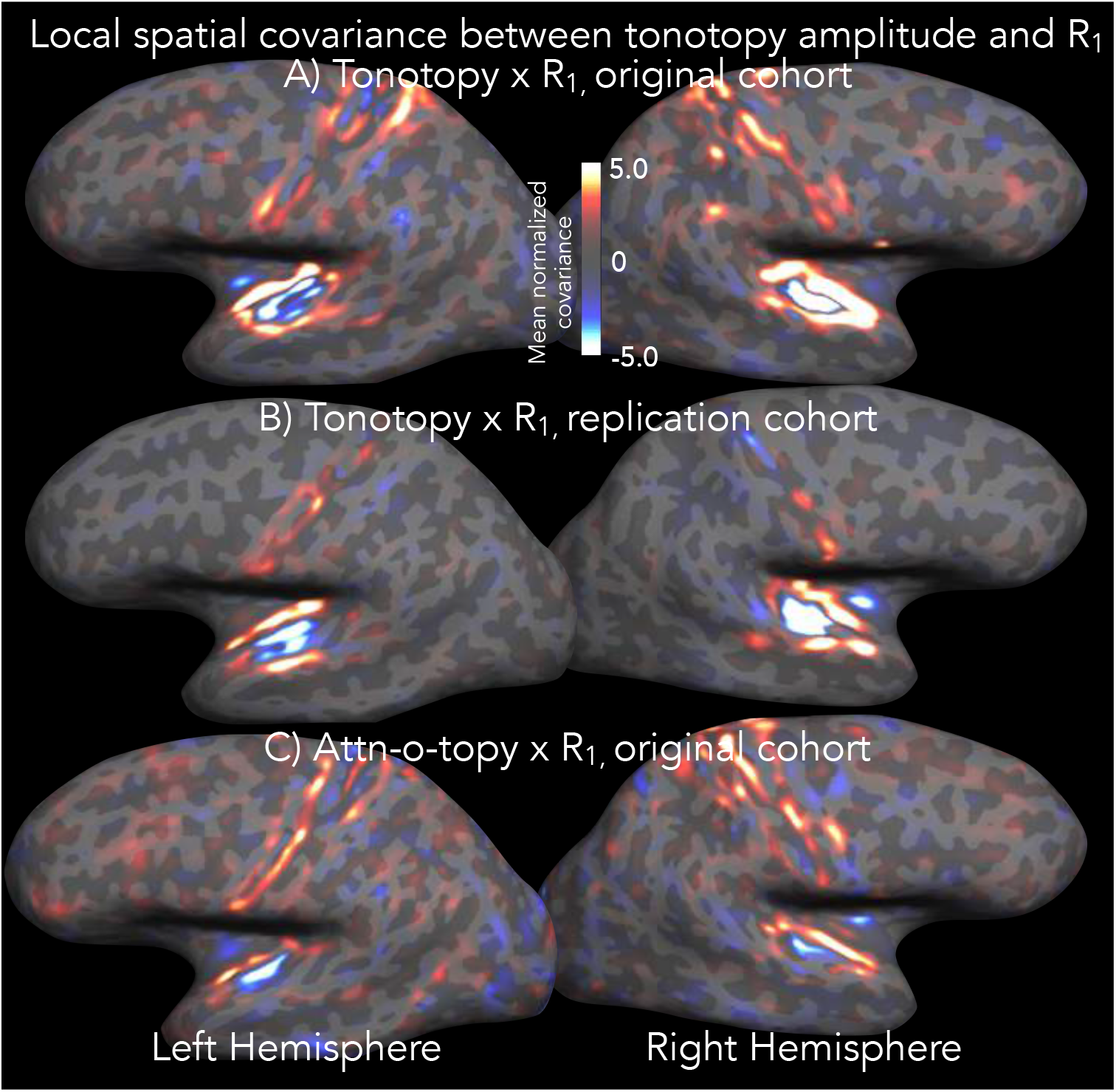
Local normalized covariance between R_1_ values and tonotopic and attn-tono response amplitude. The heatscale value at each vertex represents the normalized spatial covariance within a 4mm (2D) radius between R_1_ and the amplitude of the tonotopic or attn-tono signal (e.g., the amplitude of the Fourier component at the stimulus frequency of 8 cycles/run). **(A)** The cross-subject (N=8) cortical-surface-based average normalized covariance between R_1_ and tonotopic amplitude. **(B)** The R_1_ versus tonotopy normalized covariance in an independent cohort (N=6), using data previously acquired with a different tonotopy protocol (bandpass-filter-swept non-linguistic vocalizations) and on a different scanner (Siemens 3T Trio); full protocol as described in Dick et al., (2012)). **(C)** The average normalized covariance between R_1_ and attn-tono amplitude in the current cohort.

Here, we asked whether the change in the degree to which cortex showed a strong frequency-band preference (namely, the amplitude of the phase-encoded to not opicor attention-o-tonotopic signal) spatially corresponded with changes in myelination as assessed by quantitative R_1_ (within a 4 mm radius disk that roved across the entire cortical surface). The cross-subject-average normalized covariance map in Fig. 7a shows that there is a shared local gradient in tonotopic amplitude and R_1_ along the entire inferior circular sulcus and the anterior part of the superior temporal gyrus, where tonotopic amplitude and R_1_ drop in tandem over a narrow band of cortex. There is also negative local spatial covariance between tonotopic amplitude and R_1_ within the center of auditory cortex, where tonotopic amplitude increases but R_1_ remains relatively stable. (There is also some tonotopic/R_1_ spatial covariance within and around the central sulcus; these regions showed considerably less overall amplitude in tonotopic response, but one that spatially covaries with changes in R_1_).

To test the replicability of this novel tonotopy-versus-R_1_ searchlight cross-correlation, we reanalyzed R_1_ and tonotopy data from a previous study (Dick et al., 2012) that used a different tonotopic stimulus (bandpass-filter-swept non-linguistic vocalizations) and a slightly different multiparameter mapping protocol. Despite these methodological differences, we found a very similar pattern of tonotopic/R_1_ positive local spatial covariance within the circular sulcus and along the lateral STG, with negative spatial covariance again in the center of auditory cortex (Fig. 7b). The shared and relatively steep anterolateral and medial gradients in putative myelination and degree of frequency specificity - observed in two independently-acquired datasets - suggests a shared functional and myeloarchitectonic border, possibly similar in character to those reported recently relating resting state, standard task activation, and T1/T2-derived myelination estimates across cortex (Glasser et al., 2016; Kuehn et al., 2017).

As seen in Fig. 7c, the spatial relationship between local R_1_ and *attn-tono* amplitude changes is much less clear. Here, there is a weak relationship within and around auditory cortex that is only observed within the circular sulcus (particularly in the right hemisphere). There are also stripes of spatial covariation along the banks of the central sulcus, although not closely aligned with the pattern observed with the tonotopy versus R_1_ covariance maps. Although very preliminary, these results suggest that changes in the degree of spectral attentional modulation in auditory cortex are not strongly linked to the underlying myeloarchitecture, and stands in contrast to the consistent spatial association in lateral and medial auditory cortex between local changes in R_1_ and the strength of stimulus-driven frequency response preference.

### Summary

Everyday listening ordinarily takes place in rich soundscapes within multiple, simultaneous sound sources contributing to the overlapping mix of sound waves that arrives at the ears. Auditory attention is crucial to sorting out the mix. Listeners direct attentional focus to a sound source, or even to specific acoustic dimensions within a single sound source, to zero in on auditory information that is diagnostic in guiding behavior.

We asked how endogenous attention directed to specific acoustic frequency bands modulates human auditory cortical activity. Using high-resolution quantitative MRI and a novel fMRI paradigm for driving sustained selective attention within specific frequency bands, we established effects of spectrally-specific attention in myeloarchitectonically-estimated human auditory core. These effects extend across the majority of tonotopically-mapped auditory cortex, and are apparent in individual listeners. Sensory-driven best-frequency tonotopic maps align with attentionally-driven maps across much of the temporal plane, with poor concordance in non-auditory areas. Individual tonotopic and attention-o-tonontopic maps show correlated idiosyncracies. The frequency bands that evoke the least BOLD response from input and from attention also exhibit close spatial correspondence. There is greater activation across most of auditory cortex when best frequency is attended, versus ignored. Finally, there is local spatial correspondence in multiple auditory regions between the degree of R_1_-estimated myelination, and the strength of the frequency-band-selective fMRI response for tonotopic stimuli.

## Discussion

### Human auditory core exhibits attentionally-driven tonotopic organization

Previous findings showed similar stimulus - and attentionally-driven frequency preference in and around Heschl’s gyrus, a macroanatomical landmark associated with primary auditory areas (Da Costa et al., 2013; Riecke et al., 2016) Here, we demonstrate that, within quantitative-R_1_-defined primary auditory areas, the attentionally-driven tonotopy in each hemisphere is very similar to the detailed tonotopic maps in the same subjects. As shown by comparison maps across the acoustically-identical stepped and randomized attention-o-tonotopy conditions (Figs. 2 and 4), the alignment between tonotopic and attention maps depends on allocation of attention to the cued frequency band, not perceptual interference or other stimulus-driven effects. The fact that there is considerable, high-level attentional modulation within primary auditory areas is interesting given previous results suggesting more limited attentional topographic modulation in primary auditory (Atiani et al., 2014) and visual (Saygin and Sereno, 2008) cortex, compared to more robust attentional modulation in areas immediately adjacent to primary ones.

### Attentionally-driven tonotopic organization extends across much of auditory cortex

We also find strong evidence for tonotopically-mapped spectrally-directed attention in much of auditory cortex, particularly along the lateral superior temporal gyrus (potentially analogous to lateral auditory belt and parabelt regions in macaque (Hackett, 2007)). In addition to the concordance in and around auditory core (as defined by quantitative MR), the most consistent group-level alignment of these maps lies lateral to auditory core, with each map characterized by three higher-to-lower best-frequency-band traversals, moving from posterior to anterior roughly along the superior temporal gyrus.

This pattern suggests a cross-species parallel to results reported in ferret (Atiani et al., 2014), where task-evoked attentional modulation of frequency-tuned neurons is particularly strong in non-primary (dPEG) tonotopically mapped auditory areas in ferret. In this regard, the stimulus complexity, variability, and memory demands of the current task to a may have helped to drive attentional response in these more lateral and anterior areas. Our results are consistent with a human fMRI comparison of cross-modal attentional effects (Petkov et al., 2004), which showed greater activation in lateral auditory regions when attention was directed to a demanding auditory repetition detection task than when the same sounds were played as subjects performed a demanding visual detection task. However, they differ from these studies to some degree in that attentionally-driven tonotopic modulation in auditory core was also robust (similar to cross-modal attention studies in macaque A1 (O’Connell et al., 2014) and primary auditory areas (De Martino et al., 2015a)), and did not differ significantly from that in lateral belt.

There was good correspondence between the voxel-wise best frequency-band for tonotopy and attention-tonotopy in individual listeners. Like several prior studies (data and review in (Humphries et al., 2010; Moerel et al., 2014; Saenz and Langers, 2014; Brewer and Barton, 2016; Leaver and Rauschecker, 2016)), we observed quite substantial variation in the detailed topography of tonotopy across individuals (but cf. Ahveninen et al., (2016)). It is especially noteworthy that attention-tonotopy recapitulated these topographic idiosyncrasies (as observed in the concordance analyses, Fig. 4b and 5b).

It is intriguing that there was a systematic frequency-band-associated topography not only of best frequency but also of dis-preferred frequency and, also, that the frequency-selective attenuation of BOLD gain relative to other frequencies can be recapitulated by selective attention to that frequency band in the presence of other spectral information. One could speculate that this map structure might be a population-level reflection of an ‘inhibitory surround’ structure observed in some electrophysiology studies (Calford and Semple, 1995; Sutter et al., 1999) but cf Wehr and Zador, (2003), with the frequency band driving the least BOLD response corresponding to the deepest trough in an asymmetric surround -- an effect that could drive the very similar tonotopic and attn-tono graded frequency response preferences revealed in the multiple ROI analysis (Fig. 4c).

Here, the average frequency response profile evoked by the single-band tonotopic stimuli was recapitulated by attention to the same frequency bands in the context of distractors. Prior human neuroimaging research has been consistent with the possibility that the shape of the frequency response in and around Heschl’s gyrus is attentionally-modulated in a bandpass manner that relies on amplification rather than attenuation (Riecke et al., 2016). Based on results from a larger number of spectral bands, the current findings suggest that, at least at a more macroscopic scale, spectrally-directed attention modulates cortical activity in a more graded fashion, with the shape of the attentional response to both preferred and less-preferred frequency bands similar to that evoked by stimulus alone - a contention supported by the alignment of the ‘Loser-Takes-All’ tonotopic and attn-tono maps. That is, the frequency band that drives the smallest fMRI response when presented alone is also the frequency band that elicits the least activation when attended in the presence of a distractor. A better understanding of the mechanisms underlying these maps will require more fine-grained characterization of frequency-directed attentional modulation, preferably at very high spectral and temporal resolution (Moerel et al., 2013; Lutti et al., 2014; Moerel et al., 2014; Ahveninen et al., 2016) that might also help to unveil cortical-depth-specific attentional effects (De Martino et al., 2015a). In particular, it will be important to see whether different fMRI tasks - using more complex naturalistic sounds, or more or less abstract cues to frequency - can mimic the task-, valence-and context-dependent effects observed in non-human animal cortical auditory receptive fields, where the character of the ‘contrast-enhancing’ modulations differs markedly with experimental manipulation (Fritz et al., 2005; 2007a; 2007b; David et al., 2012; Atiani et al., 2014; Kuchibhotla et al., 2017). (It is worth noting that task-related modulation of frequency-selective attentional effects has long been of interest in human auditory psychophysics (Greenberg, 1968; Scharf et al., 1987; Scharf, 1989; Moore et al., 1996; Green and McKeown, 2001)).

### There is correspondence between local change in R_1_-estimated myelination and the strength of fMRI-assessed relative frequency selectivity

We found that the change in the degree to which a small (4 mm radius) patch of cortex shows strong frequency preferences in tonotopy was positively spatially correlated with its degree of myelination as estimated by R_1_. The strength of the correlation was anatomically specific, marking the medial border of auditory cortex (within the circular sulcus) and revealing a potential anatomical index of ‘processing style’ (from more to less tonotopically mapped) along anterolateral superior temporal gyrus. We found this pattern to hold true in the data from the current study as well as in an independent cohort scanned with quite different tonotopic stimuli and with multiparameter maps acquired on a different scanner (Fig. 7c). Although there was a relatively reliable pattern of R_1_-tonotopy correspondence at a group level, there was some notable individual variation in local shared R_1_/ tonotopy gradients relative to gyral anatomy. Thus, these patterns may be more useful than curvature for establishing areal borders on an individual subject basis, particularly when there is no obvious sharp change in a single measure (for discussion see also (Glasser et al., 2016)). Such work holds promise for generating novel hypotheses for more intensively characterized species like mouse, ferret, or marmoset, particularly in tandem with imaging techniques that that can cover multiple cortical areas simultaneously.

### Future Directions

In the current study, we limited our investigation to broadly defined auditory cortex, where there was good evidence for systematic tonotopic representation from a number of previous studies (Talavage and Edmister, 2004; Hackett, 2007; Moerel et al., 2013; 2014; Saenz and Langers, 2014; Leaver and Rauschecker, 2016). In future research it will be informative to examine interactions with several frontal regions whose potential analogues are known to have direct feedforward and feedback connections in macaque monkeys (Romanski and Goldman-Rakic, 2002), and where in ferret there are clear modulatory influences on primary and non-primary auditory cortex during learning (Atiani et al., 2014; Shamma and Fritz, 2014). Similar to recent work in vision (Klein et al., 2014; Puckett and DeYoe, 2015), it will also be useful to establish the shape of the attentional population receptive field, and how this varies across auditory areas and relates to stimulus-driven auditory population receptive field size (Thomas et al., 2015). Finally, following on our own pilot work, it will be exciting to explore whether higher-level auditory regionalization may follow along some of the ‘fault lines’ revealed by shared local tonotopic and myelin gradients, and whether or not more sophisticated and finegrained spectral attentional manipulations may reveal a relationship between the degree of attentional malleability and underlying cortical architecture and circuitry.

## Acknowledgements

Research supported by Rothberg Research Award in Human Brain Imaging of Carnegie Mellon University. M.L. was supported by NIH T90DA022761 and T32GM081760. Thanks to Scott Kurdilla and Debbie Viszlay (SIBR) for imaging support, Antoine Lutti and Nikolaus Weiskopf for physics expertise and generosity with porting the MPM protocol to the SIBR Verio, and the developers of FSL, AFNI, and FreeSurfer for their scientific work and software. Finally, we are grateful to Marlene Behrmann, Jenny Bizley, Maria Chait, Tim Griffiths, Jen Linden, Catherine Perrodin, Chris Petkov, Lars Riecke, Sam Schwarzkopf, Jeremy Skipper, Ediz Sohoglu, Adam Tierney, and three anonymous reviewers of a previous version of the manuscript for extremely useful suggestions and feedback.

